# Flexible route planning and rapid structure learning by mice in complex environments

**DOI:** 10.64898/2026.06.02.729586

**Authors:** Michael Pereira, Beatriz S. Godinho, Christian K. Machens, Rui M. Costa, Thomas Akam

## Abstract

Action selection using predictive models of the environment plays a fundamental role in human and animal behaviour, yet is poorly understood at circuit and algorithmic levels. Spatial navigation is an attractive domain for characterising how world models guide action selection. However spatial behaviours are shaped by multiple control systems including habits, vector-navigation using a Euclidean model of spatial relationships, and route planning using models of environment structure. Understanding how world models support navigation requires assays that dissociate control systems and decorrelate behavioural variables, while generating large datasets that allow precise quantification of brain-behaviour relationships. Here we developed and computationally optimised a behavioural assay to quantify flexible navigation using knowledge of environment structure. Mice navigated to visually cued goals in complex mazes, with randomised start and goal locations on each trial, generating thousands of non-repetitive goal-directed navigation trajectories. They navigated efficiently - strongly favouring options on the shortest path-to-goal, and learnt rapidly - demonstrating knowledge of maze structure from their first sessions in new environments. We anticipate the assay will be useful for characterising how world models support flexible behaviour.

## Introduction

Humans and other animals show remarkable behavioural flexibility, rapidly adapting to changes in the external environment and internal needs and goals. This is thought to be enabled by planning mechanisms that use predictive models of the environment to evaluate and select actions (Dolan and Dayan, 2013; Mattar and Daw, 2026). Despite the central importance of planning, and evidence that it is dysregulated in psychiatric illness (Gillan et al., 2016; Nour et al., 2021; Voon et al., 2015), its implementation by brain circuits is poorly understood.

A central challenge in charactering planning mechanisms is that in many situations, behaviour consistent with planning may be generated by other strategies that coexist in the brain (Balleine and Dickinson, 1998; Dickinson, 1985; Dolan and Dayan, 2013). These are thought to include model-free reinforcement learning (RL) - which learns long-run reward associated with different actions through trial and error using reward prediction errors (Daw et al., 2005; Schultz et al., 1997), value-free mechanisms which repeat actions that have previously been selected in the same situation (Greenstreet et al., 2022; Miller et al., 2019), and strategies based on working memory or state-inference (Akam et al., 2015; Collins, 2026). This multiplicity of control systems makes it challenging to isolate the contribution of planning to behaviour, or to concretely link task-related brain activity to planning computations.

An influential line of work has sought to develop behaviours that dissociate action selection using predictive models, or cognitive maps, from simpler mechanisms. Early work from Tolman and colleagues tested whether ‘latent’ learning during unrewarded exploration of a maze enabled efficient goal-directed navigation when rewards were subsequently introduced, and examined subjects’ ability to use knowledge of environment structure to exploit shortcuts or find detours around obstacles (Tolman, 1948; Tolman and Honzik, 1930). Outcome devaluation is another powerful approach developed to test for use of predictive models, building on instrumental conditioning where subjects learn to perform an action to obtain a reward, e.g. lever pressing for food pellets. After learning, the outcome is devalued by pairing it with illness or inducing sensory specific satiety, and the effect of this on action selection is tested in extinction - without further outcomes being delivered. Given moderate initial training, outcome devaluation reduces the tendency to perform instrumental actions, demonstrating that they are mediated by a prediction of the specific outcome, rather than simply knowledge that the action is valuable (Adams and Dickinson, 1981; Balleine and Dickinson, 1998). After extensive training actions can become devaluation insensitive (Dickinson, 1985), which is thought to reflect a transfer of control to habitual systems that do not utilise predictions of specific outcomes, but rather directly cache action values or preferences (Daw et al., 2005; Gremel and Costa, 2013a; Miller et al., 2019).

The development of well controlled behavioural assays has enabled causal brain manipulations to identify structures necessary for goal-directed action (Bradfield et al., 2015, 2020; Corbit and Balleine, 2003; Gremel and Costa, 2013b; Yin et al., 2005). However, it has proved challenging to leverage these tasks to characterise planning mechanisms at the circuit and algorithmic level. This is in part because many tasks designed to dissociate model-based action selection from other mechanisms provide few informative trials, as the key test must be performed in extinction, or only the first trials following a behavioural manipulation provide a clean test. To address this limitation, multi-step decision tasks were developed which aim to dissociate control systems in the context of large datasets. Subjects traverse a small decision tree on each trial, with changes in reward and/or transition probabilities necessitating ongoing learning. These have been productively used in human neuroimaging to identify signatures of model-based decision variables and associated computations (Daw et al., 2011; Gläscher et al., 2010; Huang et al., 2020), and various groups including ourselves have adapted them for animal models (Akam et al., 2021; Miller et al., 2017; Miranda et al., 2020). However, a concern with animal versions is that given the extensive training typically needed, subjects may learn to recognise the small set of different task configurations and adopt habitual mappings from these to appropriate actions. Computational modelling indicates that such strategies, formalised as a combination of state inference with model-free RL, can in principle generate behaviour that closely resembles model-based planning (Akam et al., 2015; Blanco-Pozo et al., 2024), and we have observed direct evidence for inference-based strategies in mice performing a two-step decision task (Blanco-Pozo et al., 2024).

There therefore remains a shortage of behavioural assays that are well suited to circuit-level interrogation of planning mechanisms in animal models. Spatial behaviour in rodents is a promising approach, due to its strong ethological relevance, prior work suggesting that rodents use cognitive maps to navigate, and our detailed knowledge of spatial representations (Moser et al., 2008). Spatial representations are also recruited for non-spatial tasks (Aronov et al., 2017; Behrens et al., 2018; Constantinescu et al., 2016), suggesting that understanding spatial planning may shed light on model-based action selection more broadly. Recordings in spatial tasks have greatly advanced our knowledge of the representations and dynamics that occur during reward-guided behaviour (El-Gaby et al., 2024; Ito et al., 2015; Johnson and Redish, 2007; Kay et al., 2020; Krausz et al., 2023; Pfeiffer and Foster, 2013; Sul et al., 2010; Widloski and Foster, 2022). However, laboratory spatial tasks are often repetitive, with the same small set of trajectories traversed repeatedly across trials. This makes it challenging to dissociate cognitive-map-based action selection from habitual mechanisms, and induces strong correlations among different behavioural and decision variables. For example, navigational goals are often confounded with latent task states, and spatial location with position in the sequence of trial events - complicating the interpretation of task-related brain activity.

Here we developed and optimised a behavioural assay which aimed to dissociate navigation strategies and decorrelate behavioural variables in the context of large datasets. Mice were trained to navigate to visually cued goals in complex mazes, with changing start and goal locations on each trial to minimise the utility of habitual strategies. Maze environments were numerically optimised to dissociate vector navigation - i.e. moving towards the goal in Euclidean space, from structure based navigation - moving towards the goal along the shortest available path. We acquired a large dataset of 59651 navigation trajectories including 282855 choice points. Subjects navigated efficiently, strongly favouring options on the shortest path to goal. Computational modelling confirmed a strong influence of environment structure on choices, with a weaker influence of vector navigation. Learning was rapid, with knowledge of environment structure apparent from the first session on new maze configurations. We anticipate the assay will be useful for characterising brain mechanisms of flexible behaviour in complex environments.

## Results

### Grid-maze apparatus

We aimed to develop a spatial navigation task which could dissociate the contribution of cognitive-map based navigation using a model of environment structure, from both habitual route selection and vector-navigation. These considerations suggested two design choices. First, to dissociate structure-based from vector-based navigation, the environment should have a complex topology, such that the shortest path to goal would often require moving away from the goal in Euclidean space (Figure 1a). Second, to minimise the utility of inflexible habitual strategies, start and goal locations should change from trial-to-trial.

**Figure 1:**
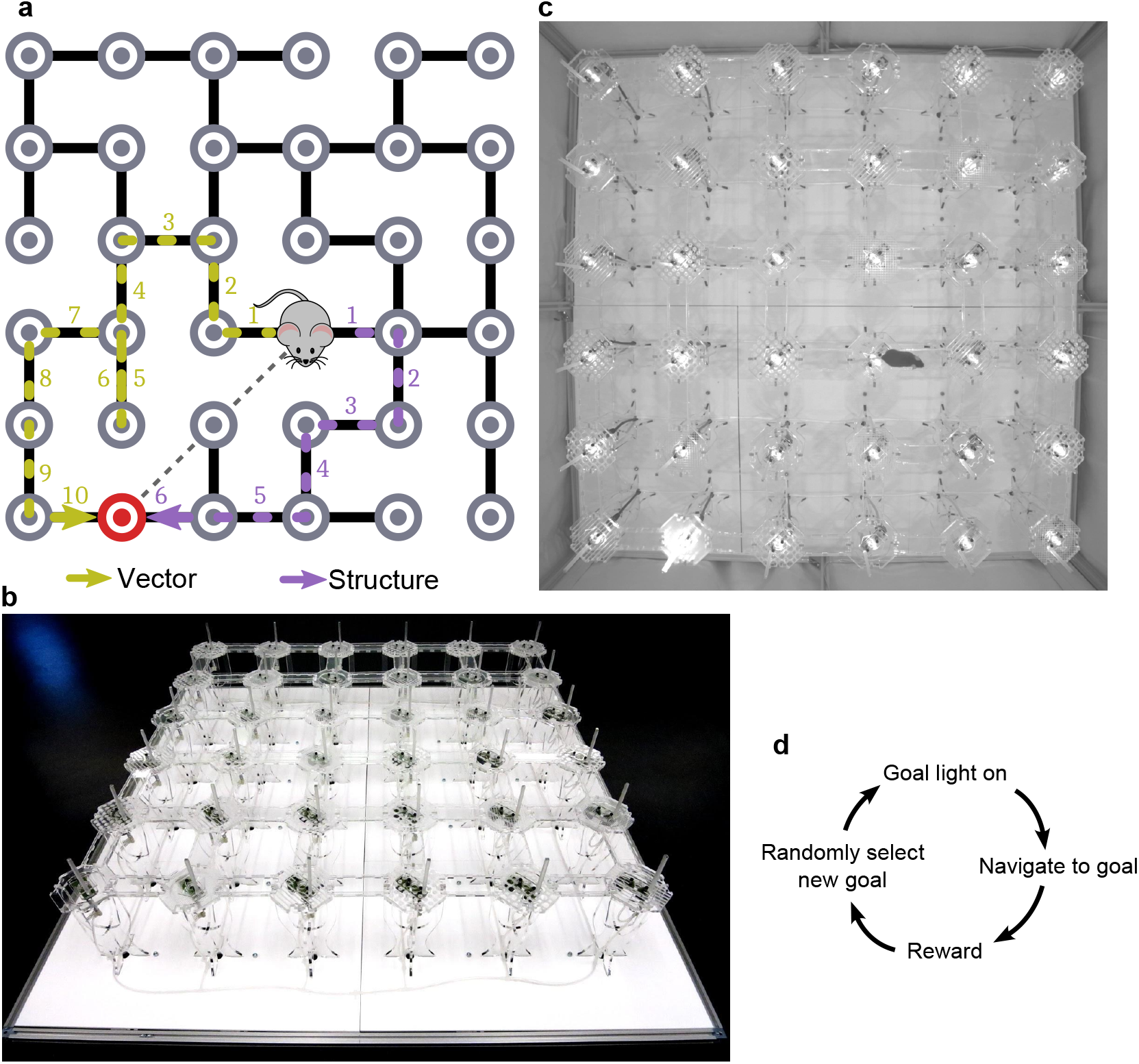
Grid-maze apparatus and route planning task: **a)** A complex maze structure creates situations where vector navigation - i.e. moving in the direction of the goal in Euclidean space (olive), favours different paths from optimal structure-based navigation - i.e. moving towards the goal along the shortest path (purple). **b)** Grid-maze apparatus showing 6×6 grid of towers, each with a reward port and stimulus LED, interconnected by removable walkways. **c)** View from overhead camera used to track animal’s position. The illuminated goal can be seen in the bottom left of the maze. **d)** Trial structure of route planning task.

To implement this we developed a behavioural apparatus we term the Grid-maze, consisting of a 6×6 grid of towers, which could be interconnected by removable walkways to form complex elevated maze layouts (Figure 1b). To allow for flexible trial-by-trial changes in goal location, each tower could be visually cued using an LED, and had a floor-mounted nose-poke port where water rewards could be delivered. Mouse position on the maze was tracked using an overhead camera (Figure 1c). The original apparatus used in this study was controlled by Arduino-based electronics, though ongoing work with the assay uses a subsequently developed version of the hardware controlled using pyControl (Akam et al., 2022), with design files available on Github.

### Route Planning Task

The behavioural assay, which we term the route planning task, consisted of a simple repeating trial structure (Figure 1d). A randomly chosen tower was selected as the goal location and cued using the tower’s LED. The subject navigated to the cued goal and poked the reward port to obtain a water reward. Following a short inter-trial-interval (ITI), another randomly chosen goal location was cued, starting the next trial.

We first piloted the task in a cohort of 8 mice to establish a training protocol and perform basic validation of the task design and apparatus. To facilitate initial learning about the cue-reward association we pre-trained the mice on a small 3×3 section of the maze, screened off from the rest of the apparatus, to increase the likelihood of the animal discovering the rewarded goal location by chance. During pre-training, subjects experienced 1 session of 40 minute duration per day, and over 11 days they increased the number of trials completed per day from 4.38 ± 2.50 to 72.88 ± 21.07 (mean ± SD; Figure S1a). They were then transferred to the 6×6 maze apparatus, using the maze layout shown in Figure 1a.

Over 9 days of training on the complex maze, subjects increased the number of trials completed per session (Figure 2a; repeated-measures ANOVA, *F*_8,40_ = 11.64, *p* =< 0.001) and learned to take more efficient routes to the goal, as indicated by a decrease in the number of excess steps per trial (Figure 2b; *F*_8,40_ = 9.07, *p* =< 0.001). To assess whether subjects were primarily using vector- or structure-based navigation we examined choice points where vector and structure disagreed, and evaluated the fraction of optimal structure-based choices (Figure 2c). To minimise the influence of within-trial adaptation following errors we considered only the first visit to each node on a given trial. The fraction of optimal choices increased over training (*F*_8,40_ = 3.60, *p* = 0.003) and was significantly above chance level on the final training day (one-sample *t*-test, *t*_7_ = 5.00, *p* = 0.002), indicating that as subjects gained experience on the maze they relied more on structure than vector-based navigation.

**Figure 2:**
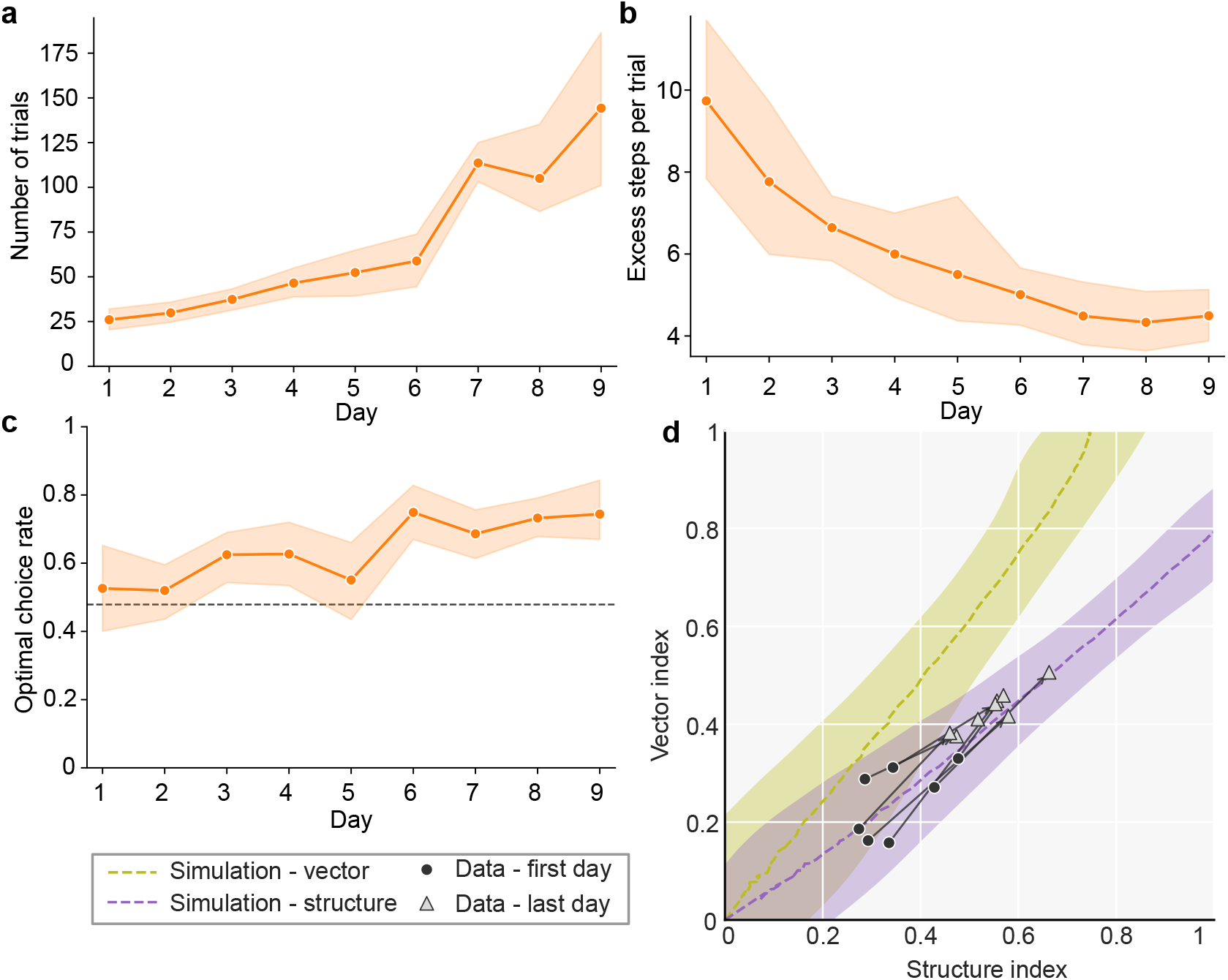
Behaviour in pilot experiment: **a)** Number of trials completed per day over 9 days of training on the 6×6 maze. **b)** Number of excess steps per trial over training, defined as the difference between the actual path length taken by the subject and the shortest path to the goal. **c)** Fraction of optimal choices at decision points where optimal structure-based navigation (reduce shortest path distance to goal) and vector-based navigation (move in direction of goal) disagreed. Dashed line indicates chance level. Plots **a**-**c** show cross-subject mean and 95% confidence interval, n=8. **d)** Subject’s behaviour on first day (circles) and last day (triangles), plotted in a space defined by vector and structure navigation indices. Each index was defined as the fraction of steps where the option favoured by that strategy was taken, normalised such that always following the strategy yields a value of 1 and a random walk yields a value of 0. Dashed lines show mean and shaded areas 95% kernel-density contour for data simulated from a structure-based (purple) and a vector-based (olive) navigation strategy at different levels of choice stochasticity.

To provide a complementary picture of the contribution of structure- and vector-based navigation strategies, and quantify how well differentiated they were given the maze layout, we defined a ‘vector-index’ and a ‘structure-index’ each indicating the fraction of choices at decision points that were consistent with the corresponding strategy, normalised such that a value of 1 corresponded to always following the strategy and a value of 0 corresponded to a random walk (see methods). Again, we considered only the choices made in the first visit to each node on a given trial. We simulated behaviour from each strategy (vector, structure) with different levels of choice stochasticity, and plotted both the subjects behaviour and the simulated data in the space defined by the two indices (Figure 2d). Behaviour on the first day was closer to the origin, indicating a more stochastic, less goal-directed strategy, which may reflect exploration of the environment competing with goal-directed navigation. By the last day, subjects’ choices moved further from the origin (paired *t*-test on distance from origin, *t*_7_ = 9.62, *p* =< 0.001), indicating more goal-directed behaviour, and were more consistent with a structure-than a vector-based strategy (mean difference in strategy index = 0.12; one-sample *t*-test, *t*_7_ = 11.32, *p* =< 0.001). These data show mice can learn to navigate complex environments to reach visually cued goals, and the resulting behaviour reflects knowledge of the environment structure.

### Maze structure optimisation

The maze layout used in the first experiment (Figure 1a) was hand designed, and hence is unlikely to be optimal for dissociating structure- and vector-based navigation. Indeed, simulations from structure- and vector-based agents were not widely separated in the space defined by metrics of structure- and vector-consistent choices (Figure 2d), reflecting the fact that for most current and goal location pairs, vector and structure-based navigation mandated the same choice. We therefore numerically optimised the maze structure to increase discriminability between strategies.

We randomly generated a large ensemble of connected maze layouts (1.7 × 10^5^ mazes) and first considered the *‘fraction of informative states’* as a measure of how well a maze discriminated structure-from vector-based navigation, defined as the fraction of current and goal location pairs for which the two strategies mandated different choices. Mazes that scored highest on this criterion tended to comprise an extended linear structure folded into the 6×6 grid (see *Maze A* in Figure 3b and additional examples in Figure S2). Navigation on such mazes is an interesting test case, but may not be representative of complex environment navigation in general, as the decision is largely reduced to deciding which way to move along the linear structure. It is therefore desirable to identify mazes where strategy discrimination arises from a set of smaller structural features.

**Figure 3:**
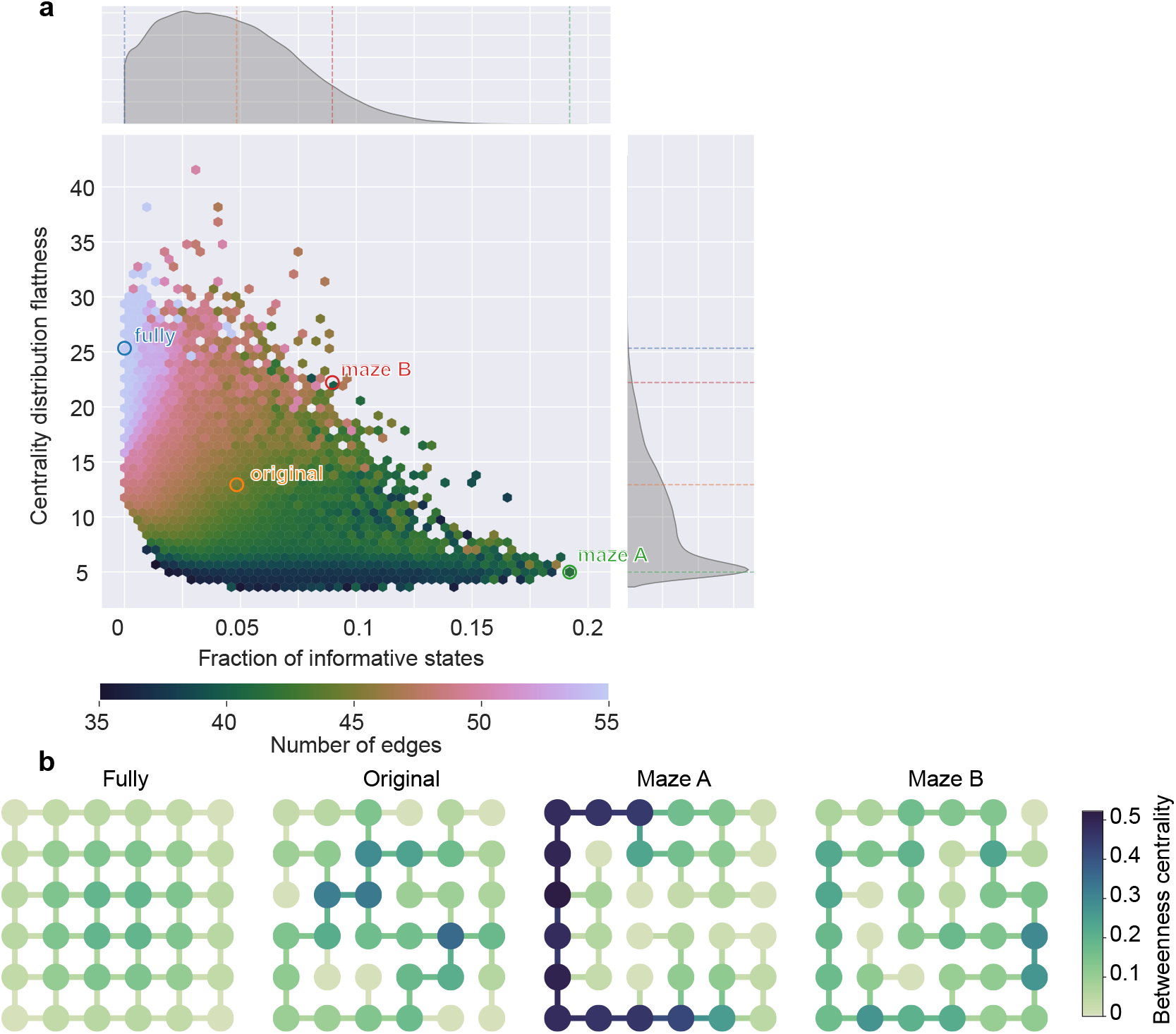
Maze structure optimisation: **a)** Distribution of mazes generated during maze optimisation in a space defined by the fraction of states in which vector and structure based navigation favour different choices, and a measure of how flat the distribution of betweenness centrality values is across maze locations. **b)** The 4 mazes used in experiment 2, coloured by betweenness centrality. The position of these mazes in the optimisation space is indicated by labelled points in **a**.

To obtain a more diverse family of mazes we evaluated each generated maze using two metrics: the fraction of informative states considered previously and a measure of how flat the distribution of betweenness-centrality values was across maze locations (see methods). Betweenness centrality is a graph-theoretic measure defined for every node and edge, whose value is the fraction of shortest paths between all pairs of nodes that the given node/edge is on, such that locations with high betweenness centrality lie on many shortest paths. We reasoned that for mazes dominated by a single linear feature, betweenness centrality would be very high at the centre of the line and very low at the ends, causing a wide distribution across locations (Figure 3b). By contrast, for mazes where many smaller features generate complexity, we expected the distribution of betweenness centrality values to be flatter. As expected, the sampled maze layouts showed a trade-off between the fraction of informative states and the flatness of the centrality distribution (Figure 3a). For use in our second experiment we selected two mazes (*Maze A* and *Maze B*, Figure 3b) that lay on the empirical Pareto-optimal front, meaning that no sampled maze improved one of the two metrics without worsening the other. The two mazes therefore represented different trade-offs between these metrics. Our original maze was far from this front, indicating it was not optimal with respect to the combination of these two criteria (Figure 3a).

### Mouse behaviour across maze topologies

To characterise mouse navigation behaviour across a set of different maze topologies, we trained a new cohort of 8 mice sequentially on a series of 4 different maze structures (Figure 3b). These were; a fully connected maze, the original hand-designed maze used in the first cohort, *Maze A* - the optimised maze which best discriminated vector and structure-based navigation strategies, and *Maze B* - another optimised maze which represented a trade-off between the fraction of informative states and a flat distribution of betweenness centralities. We pre-trained the mice to understand the cue-reward association in a separate small 3×2 maze (Figure S1c,d), before transferring them to the 6×6 maze.

The number of trials completed per session increased with experience within each maze, except the final maze where the number of trials remained at a stable, high level throughout (Figure 4a; fully connected, *F*_8,56_ = 15.20, *p* =< 0.001; original, *F*_9,63_ = 2.96, *p* = 0.005; Maze A, *F*_11,77_ = 11.72, *p* =< 0.001; Maze B, *F*_9,63_ = 1.38, *p* = 0.216). The number of excess steps per trial decreased with experience on each maze as mice learned to navigate them efficiently (Figure 4b; all *F >* 11.9, all *P* < 0.001). This effect was strongest for *Maze A* - optimised to maximally discriminate structure from vector-based navigation, indicating that the resulting maze was challenging for subjects to learn to navigate efficiently. Excess steps decreased with experience on the fully connected maze despite vector navigation being sufficient to select optimal paths. This may reflect subjects prioritising exploration when they were first exposed to the 6×6 maze environment. Consistent with this, the learning curve for excess steps on the ‘original’ maze layout, used in both the first and second experiments, started at a lower level (less excess steps) in the second experiment, where subjects already had extensive experience with the fully connected layout, than in the first experiment when the apparatus was novel (Figure 2b).

**Figure 4:**
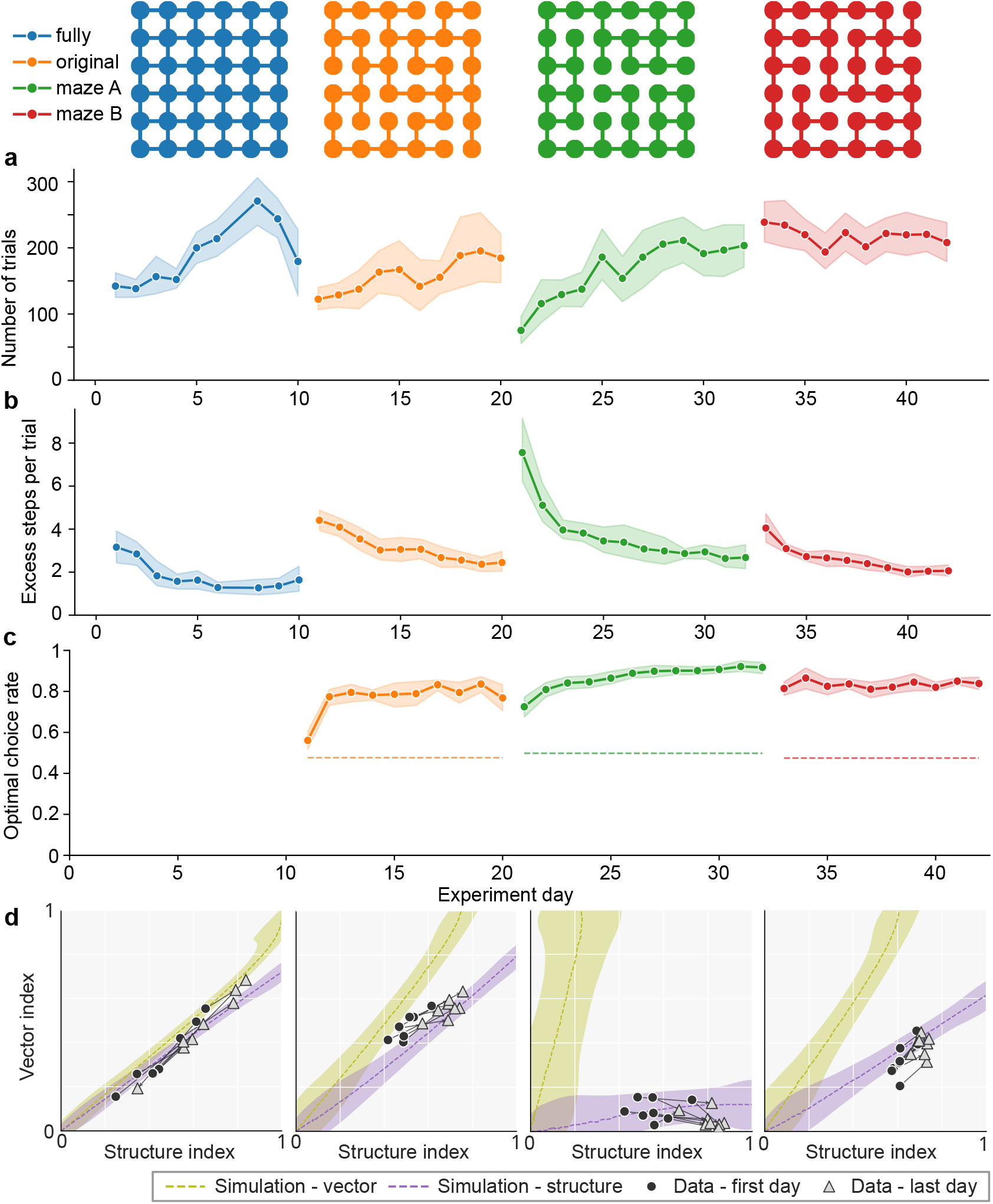
Mouse behaviour across maze topologies. **a)** Number of trials completed per experiment day across the 4 sequentially presented maze configurations, indicated by colour. **b)** Number of excess steps per trial. **c)** Rate of optimal structure-based choices at decision points where vector- and structure-based navigation disagree. Dashed line indicate maze-specific chance levels. **d)** Behaviour of each subject on each maze plotted in the strategy space used in Figure 2d.

To examine how navigation strategies changed within and across mazes we first plotted the fraction of optimal structure-based choices at first visits to nodes where vector and structure disagreed, excluding the fully connected maze where the strategies always agreed (Figure 4c). Subjects chose the optimal option slightly above chance level on the first day of the *original* maze (one-sided t-test, *t*_7_ = 3.31, *p* = 0.006), with a sharp increase on the second day to a sustained high level of optimal choices. On *Maze A* the fraction of optimal choices was well above chance level on the first day (*t*_7_ = 7.61, *p* =< 0.001), and continued to rise gradually with experience on the maze. On *Maze B* the fraction of optimal choices was at a sustained high level from the first day on the maze (*t*_7_ = 14.64, *p* =< 0.001). These data indicate very rapid learning of maze structure, with most learning happening within the first day for mazes A and B, and within the first two days for the original maze. This different learning dynamics may reflect the fact that the transition from fully connected to original maze was the first time subjects experienced a configuration change, and a complex maze layout that necessitated using knowledge of maze structure.

To further visualise the evolution of navigation strategies, we plotted subjects’ behaviour in the space defined by vector and structure strategy indices, along with behaviour simulated from single-strategy agents at different levels of choice stochasticity (Figure 4d). Simulated data from the two strategies was most well separated for *Maze A*, as expected from this maze being optimised solely to discriminate these strategies. *Maze B* - optimised to discriminate strategies while maintaining a flat betweenness centrality distribution, separated strategies more effectively than the original hand-crafted maze, consistent with the larger fraction of informative states (Figure 3a). As expected, vector and structure based navigation were not differentiated in the fully connected maze.

The first complex maze (*original*) was common to experiments 1 and 2, but the evolution of subjects’ behaviour in strategy space was somewhat different across experiments (Figure 2d, Figure 4d). Behaviour from the first day on experiment 2 was both further from the origin (i.e. less stochastic, more goal oriented) and had a stronger contribution of vector navigation. This may reflect the fact that in experiment 2 subjects had been exposed to the fully connected maze layout for 10 days, which may have both reduced exploratory drive and biased them towards a vector-navigation strategy. With experience on the original maze in experiment 2 subjects’ behaviour became more consistent with structure-based navigation. On both optimised maze layouts, behaviour from both the first and last day was consistent with simulations from a structure-based agent, with a small decrease in stochasticity with more maze experience.

These data show that complex maze navigation in this assay was primarily driven by structure-based rather than vector navigation, and that new maze structures were learnt very rapidly on first encountering a new environment.

### Mixture-of-strategies model

To further quantify the contribution of different navigational strategies to subjects’ behaviour, we developed a computational model which predicted subjects’ choices step-by-step during navigation, using a weighted combination of components capturing different influences on choice (Figure 5a). Each component assigned a scalar-valued preference to each available action (north, south, east, west) at every step. Component preferences were linearly combined using a set of weights which determined how strongly each component influenced choices. The resulting net preference determined choice probabilities via a softmax nonlinearity. The model was fit to behaviour using hierarchical Bayesian inference, which used the choice likelihood under the softmax action policy (i.e. the probability assigned by the model to subjects’ observed choices) to jointly infer the group-level distribution of component weights and approximate posterior distributions over subject-level weights (Piray et al., 2019).

**Figure 5:**
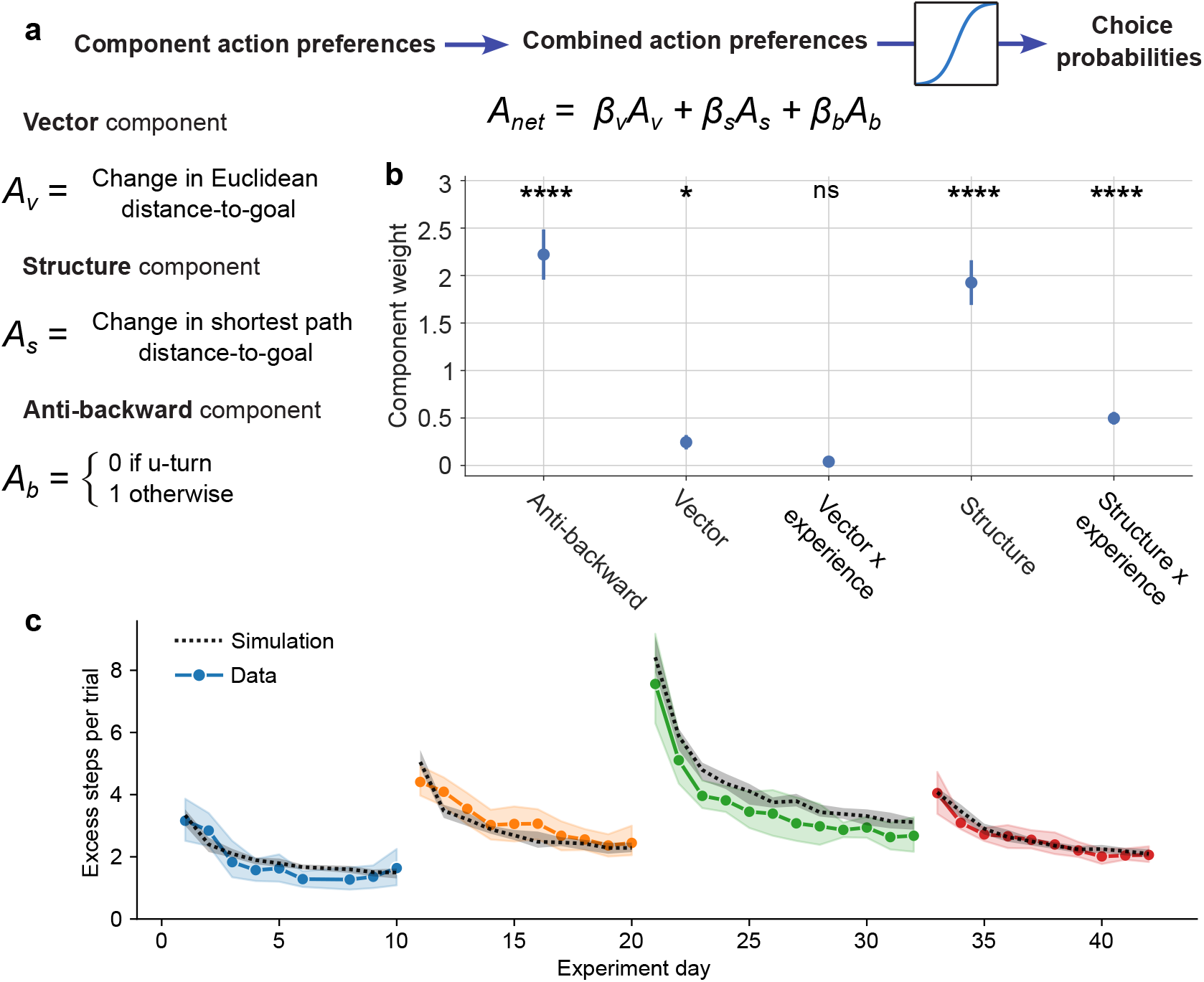
Mixture of strategies model. **a)** Diagram of model structure. The model predicted movement direction at every step during navigation using a weighted combination of components, each of which assigned a scalar preference to each available action. These were: *Vector* - move towards the goal in Euclidean space, *Structure* - move towards the goal along the shortest path. *Anti-backward* - avoid u-turns during navigation. Component action preferences were combined in a linear sum using component weights (betas) to yield a net preference, which determined choice probabilities via a softmax nonlinearity. **b)** Component weights in mixture of strategies model fit to behaviour from all four mazes used in the second experiment. For the vector and structure components, additional predictors for the interaction between the strategy and amount of experience on the current maze were included to capture learning effects. Error bars show the hierarchical error around the group posterior component weight means and significance markers indicate the results for the HBI t-test of each group parameter against zero: ns, *p* ≥ 0.05; *, *p* < 0.05; **, *p* < 0.01; ***, *p* < 0.001; ****, *p* < 0.0001. **c)** Learning curves for excess steps for simulated data from fitted model (black) and subject’s data (colour - duplicated from Figure 4b), with matched numbers of trials per session.

The components included in the model were: *Vector* - whose action preferences were given by the change in Euclidean distance to goal produced by the action; *Structure* - the change in shortest-path distance to the goal; and *Anti-backward* - which preferred not making a U-turn and returning to the previous location. The latter was included because subjects very rarely made U-turns once they had commenced navigation to the goal (Figure S3). To capture possible changes in strategy use with experience on each maze, we added two interaction components, *Vector* × *experience* and *Structure* × *experience*, in which experience was coded as *log(number of steps)*.

We fit the model to the complete dataset from the second experiment and plotted the weights for each component (Figure 5b). Weights for the *Vector* and *Structure* components were both significantly positive (hierarchical tests against zero: Vector, *t*_9_ = 3.23, *p* = 0.010; Structure, *t*_9_ = 8.18, *p* =< 0.001), indicating that both strategies influenced choice behaviour, though the weight of the Structure component was substantially larger. The *Structure* × *experience* component also had a significant positive weight (*t*_9_ = 7.45, *p* =< 0.001), indicating that the influence of structure-guided navigation increased with experience on a given maze. By contrast, the group-level loading on the *Vector* × *experience* component was not significantly different from 0 (*t*_9_ = 1.52, *p* = 0.164), providing no clear evidence that maze experience systematically changed the influence of vector navigation at the group level. There was also strong positive loading on the *Anti-backward* component (*t*_9_ = 8.39, *p* =< 0.001), consistent with subjects’ tendency not to make U-turns during navigation.

To assess the extent to which the model captured learning effects within and across mazes, we simulated data from the fitted model, with numbers of trials per session matched to subjects’ data. Simulated learning curves for excess steps closely matched that of the real subjects (Figure 5c), indicating that using strategy-experience interaction terms, with the amount of experienced quantified as *log(number of steps)*, provided a good approximation to subjects’ learning.

This behavioural modelling, in combination with the analyses of choice behaviour presented earlier, indicate that in navigating complex environments to visually cued goals, mice use knowledge of the environment structure - favouring options on the shortest path to goal, but also show a weaker influence of vector navigation.

## Discussion

We developed and computationally optimized a behavioural assay that dissociates competing navigation strategies while generating large-scale, flexible, non-repetitive behaviour, designed for circuit-level interrogation of planning mechanisms. Mice navigated a complex, reconfigurable, elevated maze to reach visually cued goals that changed on every trial. We numerically optimised maze layouts to discriminate vector from structure-based navigation, identifying a trade-off between maximising discriminability and avoiding layouts dominated by a single linear structure. We acquired a large dataset on a series of maze topologies, and quantified behaviour using analysis of choices and behavioural model fitting. Both approaches demonstrated a strong influence of structure-based navigation - choosing options that reduce the shortest path distance to goal, with the model also indicating a weaker contribution of vector-navigation. Learning was rapid, with subjects demonstrating knowledge of maze structure from their first session in a new environment. This combination of rapid learning, and flexible, efficient goal-directed navigation, is consistent with model-based action selection, though further work will be needed to conclusively determine underlying mechanisms.

Other recent studies have observed rapid learning about the structure of complex spatial environments (Cothi et al., 2022; Rosenberg et al., 2021; Widloski and Foster, 2022), likely reflecting the ethological validity of spatial behaviour. A key feature of the present task is the generation of large datasets of non-repetitive goal-directed navigation trajectories through complex environments, combined with analyses to dissociate navigation strategies. Changing start and goal locations on each trial necessitate behavioural flexibility, but critically also decorrelate behavioural variables which are often confounded in traditional spatial tasks. Specifically, the assay decorrelates position in physical space from both position in the sequence of trial events (cue-navigation-reward-ITI) and the current goal location. Further, the complex structure of the environment will decorrelate spatial representations and decision variables (e.g. distance-to-goal) that use a Euclidean representation of space from those that use a topological representation that respects the transition-structure of the environment.

Using visually cued rather than remembered goals dissociates the cognitive demands of navigating a complex environment from spatial memory for goal location, though a limitation of this design choice is that visual acuity may limit the accuracy of goal location estimates at long distances. The grid-maze apparatus is also well suited to spatial memory tasks, for example we have recently used a 3×3 grid-maze to characterise mechanisms of sequence working memory in frontal cortex (El-Gaby et al., 2024). It would be interesting to extend the route planning task to incorporate choosing among or sequencing multiple goals, introducing a hierarchical element to the task structure. Another possible extension would be to introduce local changes to the maze structure, for example adding or removing one link every session, creating an ensemble of shortcut and detour problems that could be analysed in a common framework.

Our behavioural analyses and modelling demonstrate that subjects preferentially selected options that are on the shortest path to goal, what computational mechanisms might underlie this? Model-free reinforcement learning using spatial locations as states is not sufficient, as a fixed value function or policy defined over spatial location will not adapt to trial-by-trial changes in goal location. In principle, model-free RL operating over the product space of spatial- and goal-location could solve the task, by learning separate spatial value functions for each different goal. However, we think it unlikely that this is how the brain solves the task, both on normative and empirical grounds. Normatively, such a solution is highly sample inefficient - as learning does not generalise between goals, and is inflexible in the face of changes to environment structure or introduction of new goals. The rapid emergence of structure-based navigation we observed suggest efficient mechanisms for structure learning that generalise across goals, combined with flexible use of this knowledge to reach the current goal. This is consistent with prior work on spatial learning which suggest that rodents learn spatial cognitive maps that support flexible navigation (Cothi et al., 2022; Rosenberg et al., 2021; Tolman, 1948; Tolman and Honzik, 1930). Recordings of place cells in the hippocampal formation suggest that place fields reflect the environment’s transition structure rather than Euclidean space - fields are strongly influenced by barriers and do not typically span them (Barry et al., 2006; Muller and Kubie, 1987; Skaggs and McNaughton, 1998; Widloski and Foster, 2022), providing a possible substrate for a cognitive map of environment structure (Gustafson and Daw, 2011; Muller et al., 1996; Stachenfeld et al., 2017).

The neural computations through which cognitive maps support action selection remain poorly understood. A diverse set of algorithms have been proposed, including sequential planning by simulating possible future action sequences (Mattar and Daw, 2018), representation-based methods that store long-range relationships (e.g. distances) between states (Dayan, 1993; Piray and Daw, 2021; Sagiv et al., 2025), or approaches which infer future trajectories using recurrent dynamics (Corneil and Gerstner, 2015; Jensen et al., 2025). Internally generated sequential activity in the hippocampal formation is a possible substrate for sequential planning (Johnson and Redish, 2007; Kay et al., 2020; Pfeiffer and Foster, 2013; Widloski and Foster, 2022). Other lines of evidence suggest mechanisms that avoid the need to plan sequentially by embedding long-range relationships among states in neural representations (Momennejad et al., 2017; Russek et al., 2017; Stachenfeld et al., 2017). Indeed, current data suggest that planning may itself comprise a set of mechanisms that learn different types of models, and use different computational principles, to evaluate and select actions (Mattar and Daw, 2026).

Understanding how planning works at a circuit and computational level remains an outstanding challenge for behavioural neuroscience. We hope that the behavioural assay developed in this work can contribute to uncovering these mechanisms.

## Acknowledgements

We thank Peter Dayan for useful discussions about the work, and Peter Doohan and Kristopher Jensen for comments on the manuscript. We thank Gonçalo Lopes for support in the development of the computer vision pipeline used for mouse tracking in the preliminary experiment, Hélio Rodrigues for support in assembling the behavioural box used in the pre-training phase of the main experiment. The work was supported by the Wellcome Trust (WT096193AIA, 214314/Z/18/Z, 225926/Z/22/Z to T.A.), the National Institutes of Health (5U19NS104649 to R.M.C.), the European Research Council (CoG 617142 to R.M.C), the Simons Foundation (SCGB #543009 to C.M.), and the Fundação para a Ciência e Tecnologia (SFRH/BI/52018/2012, SFRH/BD/52222/2013, to M.P., SFRH/BD/138752/2018 to B.G.).

## Methods

### Apparatus

The towers and walkways of the grid-maze were constructed from laser-cut acrylic, with a custom designed circuit-board mounted under each tower containing the IR components for the nose-poke, the LED used to visually cue goal locations, and a connector for a miniature solenoid used for reward delivery (Figure S4). The LED illuminated a frosted acrylic rod mounted vertically on the tower, providing a visual cue that could be seen across the maze. The towers were hexagonal, of size 11cm from edge-to-edge, and the walkways were 7cm long and 4.4cm wide, such that the centre-to-centre spacing between towers was 18cm and the total distance across the 6×6 grid was 1m. The walkways had low walls along each side of height 5mm to provide a secure footing. The tower tops were laser engraved with different textures to provide local spatial cues. The poke PCBs were connected to a control PCB which used an Arduino Mega 2560 to control their inputs and output programmatically. The apparatus was housed in an enclosure made from aluminium rail and covered with fabric. Back-lit extra-maze cues were mounted on the walls of the enclosure 30cm above the level of the towers.

Complete design files for an updated version of the hardware, recommended for new projects due to an improved pyControl-based (Akam et al., 2022) control system and more robust construction, are available at: https://github.com/pyControl/hardware/tree/master/GridMaze

### Subjects

All procedures were reviewed and performed in accordance with the Champalimaud Foundation Ethics Committee guidelines. 8 mice were used in the pilot experiment (Figure 2) and 8 in the main experiment (Figure 4). Mice were C57BL/6 males bred at the Champalimaud Center for the Unknown, aged 8-12 weeks at the start of the experiment. Animals were housed under a 12 hours light/dark cycle with experiments performed during the light cycle.

### Behavioural training

Mice were placed on water restriction 48 hours before the first behavioural training session, and given 1 hour *ad libitum* water access in their home cage 24 hours before the first training session. Mice received 1 training session per day of approximately 45 minute duration. On days off from behavioural training mice had 1 hour *ad libitum* water access in their home cage. On training days mice typically received all their water in the task (typically 0.5-1.25ml), but additional water was provided as required to maintain a body weight > 85% of their pre-restriction weight.

Both pilot and main experiments comprised a pre-training phase in which subjects learned the cue-reward association on a small maze layout, followed by the full task on the 6×6 maze. The rationale for pre-training on a small maze was to increase the likelihood of subjects encountering the rewarded goal location by chance before they understood the cue-reward association. For the pilot experiment pre-training was carried out in a small section of the 6×6 maze screened off from the rest of the apparatus, while for the main experiment a dedicated pre-training setup was used.

Both training stages used the same trial structure. Each trial started with a randomly selected tower being chosen as the goal location and cued using the tower LED. The LED remained illuminated until the subject poked the nose-poke port at the goal, triggering the LED to turn off and a water reward to be delivered in the nose-poke port. Pokes in non-goal ports were recorded but had no effect on the task. After an ITI, the next trial started with a new goal being randomly selected and cued. In the pilot experiment the ITI was a fixed duration (2-3 seconds) following reward delivery. In the main experiment the ITI lasted 1.5 seconds after the animal poked-out at the end of reward consumption. Goal locations were sampled without replacement until all goals had been used, at which point the sampling process was reinitialised with all goals again available for selection.

For the pilot experiment pretraining took 11 days. The the first 7 days of pretraining used a 4 node “T” shaped maze configuration, while the last 4 days used a 7 node “H” shaped configuration (Figure S1a). Over the course of of pre-training, reward sizes were reduced gradually from 32µl to 10µl as subjects learned the cue-reward association and were able to complete more trials per session.

For the main experiment, the animals were pretrained for longer (17 days) to increase the number of trials per session before transferring them to the 6×6 maze. All pretraining days used a 6 node “H” shaped maze configuration (Figure S1c). Reward sizes were reduced from 32 to 5µl over the course of pretraining.

### Analysis of behavioural performance

For analysis maze layouts were represented as undirected graphs in which towers were nodes and walkways were edges. Geodesic distance, *d*_*G*_(*i, j*), was defined as the length of the shortest path between nodes *i* and *j* on this graph, while Euclidean distance, *d*_*E*_(*i, j*), was the straight-line distance between the corresponding positions on the grid.

#### Excess steps per trial

For each trial *t*, we quantified route inefficiency as the number of steps taken in excess of the shortest available route,

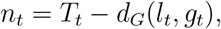

where *T*_*t*_ is the number of steps taken to reach the goal, *l*_*t*_ is the animal’s location at goal cue onset, and *g*_*t*_ is the goal location. Excess steps were averaged over trials for each subject and day; plotted lines show the cross-subject mean and 95% confidence intervals.

#### Optimal choice rate at informative choice points

We calculated the rate at which subjects took optimal choices at decision points where vector and optimal structure-based choices disagreed (Figure 2c). To isolate decisions that distinguished vector-from structure-based navigation, we analysed choice points with three or four available actions and excluded the u-turn action that would return the animal to the preceding location. To minimise the influence of within-trial adaptation following errors, we considered only the first visit to each node on each trial. For current location *l*, goal *g*, and the set of possible next locations *l*^′^ ∈ 𝒩 resulting from taking the remaining actions, we defined the preferred sets of a structure-based agent as

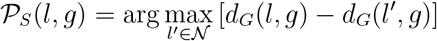

and a vector-navigation agent as

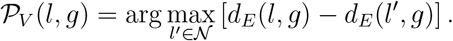

A choice point was classed as informative when these preferred sets were disjoint, 𝒫_*S*_(*l, g*) ∩ 𝒫_*V*_ (*l, g*) = ∅. Choices was counted as optimal when they belonged to 𝒫_*S*_(*l, g*). Optimal choice rate was computed for each subject and maze day as the fraction of informative choices that were optimal. Chance level for each informative choice was |𝒫_*S*_(*l, g*) |*/*|𝒪|; the maze-specific chance line shows the pooled mean of this quantity across all analysed choices in that maze.

#### Vector and structure indices

To summarise the relative influence of vector and structure-based navigation, we calculated a structure index and a vector index from the first visits to each decision point within each trial (Figure 2d, Figure 4d). At each decision point, u-turns were omitted from the set of available actions, and steps that were u-turns were not scored. A structure opportunity was any decision for which the remaining actions differed in their geodesic distance to the goal; a vector opportunity was defined analogously using Euclidean distance. For each strategy *q* ∈{*V, S*} we counted the number of opportunities *o*_*q*_, the number of choices consistent with the strategy *c*_*q*_, and the expected number of consistent choices under random choice among the available actions *r*_*q*_. The corresponding index was

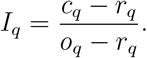

Thus an index of 0 corresponds to random choice, while an index of 1 corresponds to always choosing an action preferred by the strategy when it made a discriminable prediction. We calculated these indices for each subject on the first and last sessions for each maze layout and joined paired observations from each subject with arrows.

For comparison we computed the indices for data simulated using either a vector- or structure-based strategy with different levels of choice stochasticity, indicated by the coloured regions in the index plots. Simulated data were generated using the mixture-of-strategies model detailed below. For vector- or structure-based agents, the weight *β*_*V*_, *β*_*S*_ for the corresponding strategy was set to one of a range of 100 logarithmically spaced values from *e*^−2.5^ to *e*^2.5^ to generate a family of agents with different levels of choice stochasticity, while the weight of the other strategy was set to zero. All agents also included an anti-backward component which avoided making u-turns during navigation, as this was prominent in subjects behaviour (Figure S3), with the weight of this component set to *β*_*b*_ = 2.2 to match the group-level mean in the model fit to subjects’ data (Figure 5).

We simulated 1,000 sessions for each value of *β*_*V*_ and *β*_*S*_. To set the simulated session length for each panel, we used whichever of the empirical first- or last-day session groups contained fewer total steps, and matched the simulations to the mean total number of steps per session in that reference group. Simulated trials were capped at the 99th percentile of step counts per trial (45 steps) in the experimental data. If a simulated trial reaches this threshold without the agent reaching the goal, the agent was placed at the goal to start the next trial, to maintain coverage over locations and goals. Simulations were summarised with the same first-visit index calculation as the data, and the shaded regions show 95% kernel-density contours of the resulting simulated index distributions.

### Maze optimisation

To survey maze layouts that separated structure-based from vector navigation, we randomly generated a large ensemble of connected 6 × 6 mazes. We sampled 10,000 mazes for each of 17 edge counts (35–50 edges and 55 edges). For a requested edge count, generation began from the fully connected grid, edges were randomly removed until the target count was reached, if the resulting graph was disconnected, randomly selected present and absent edges were exchanged until all nodes were mutually reachable.

For each generated topology, we quantified the fraction of informative states using the same two deterministic decision rules defined above, but before removing backward actions. Ordered location-goal pairs (*l, g*) with *l* ≠*g* were evaluated only when *l* had at least three neighbours. A pair was labelled informative when the set of actions that maximally reduced Euclidean distance and the set of actions that maximally reduced geodesic distance were non-overlapping. The maze-level score was

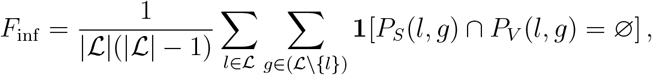

where ℒ is the set of all locations and 1 is an indicator that is zero for locations with fewer than three neighbours.

We found that optimising only for the fraction of informative states typically generated mazes dominated by a single linear structure (Figure 3b, Figure S2). To generate mazes that achieved strategy discrimination with a set of smaller structural feature, we jointly optimised for fraction of informative states and a measure of how flat the distribution of betweenness centrality values was across the maze, defined as *µ/σ*^2^, where *µ* and *σ*^2^ were the mean and variance of the node betweenness-centrality distribution. To jointly optimised these two metrics we choose mazes from the ensemble of randomly generated mazes that lay on the Pareto front, i.e. mazes for which no other maze had a higher value of both metrics.

In Figure 3 each hexagonal bin summarises generated topologies with similar values of these two metrics, and colour denotes the mean number of edges in the bin; the maze layouts used for the experiment were evaluated by the same procedure and marked accordingly.

### Model comparison and fitting

We modelled subjects’ behaviour using a mixture-of-strategies model in which a set of components, each implementing a different strategy, contributed additively to a softmax action policy. Mathematically, this formulation is equivalent to a conditional logit model from the discrete-choice literature (McFadden, 1974; Train, 2009). At each decision point the model assigned action preferences as:

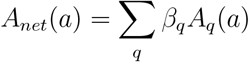

Where *A*_*q*_(*a*) is the preference for action *a* due to component *q*, and *β*_*q*_ is the associated component weight.

The net action preferences determined choice probabilities as:

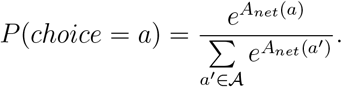

where *a*^′^ ∈ *A* are the set of available actions at the decision point.

We used three base strategy components, *vector, structure*, and *anti-backwards*. The *vector* component assigned action preferences as the change in Euclidean distance to the goal resulting from the action:

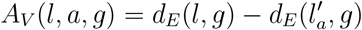

where *l* is the current location, *g* is the goal location, and the location reached after taking the action. The *structure* component assigned action preferences as the change in shortest path distance to the goal:

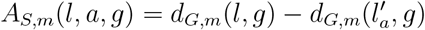

The anti-backwards component assigned a preference of 0 if the action was a u-turn and 1 otherwise. To model possible changes in strategy with experience, we incorporated components for the interaction of each strategy with the amount of experience on the current maze, with the experience moderator defined as log(maze step count + 1).

Following Gelman (2008), the vector and structure components, and experience moderator, were scaled by twice their empirical standard deviation, while the binary anti-backwards component was kept on a 0/1 scale. For the vector and structure components, which take a vector value for each choice with elements corresponding to the available options, the standard deviation *σ*_*q*_ for strategy *q* was computed as:

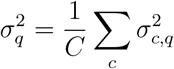

where *C* is the number of choices, and 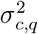 is the within-choice-set variance for strategy *q*’s action preferences at choice *c*, computed using Bessel’s correction for small sample size. Interaction terms were formed from the already-standardised input variables, rather than standardising the interaction products afterwards ((Gelman, 2008)).

We compared a full model containing the three base components and all three experience interactions against six nested variants in which different component-experience interactions were removed (Table S1). Group-level inference and Bayesian model comparison were performed with hierarchical Bayesian inference (HBI) using our Python implementation of the HBI algorithm from Piray et al. (2019), available at https://github.com/michaelfsp/pycbm. The remaining HBI hyperparameters were set to the values recommended by Piray et al. (2019), corresponding to a prior weight of one experimental sample. The model-comparison summary in Table S1 reports estimated population model frequency and protected exceedance probability for each candidate model. For the winning model, plotted parameter estimates in Figure 5b are the inferred group means with hierarchical error bars; significance labels were obtained from hierarchical tests of each group parameter against zero.

#### Posterior predictive simulations

To assess whether the fitted model reproduced the temporal dynamics of learning, we generated posterior predictive trajectories from the winning model, preserving the empirical trial schedule for each subject. For each of 16 replicate draw sets, we drew one parameter vector for each subject from that subjects’ approximate posterior distribution under the winning model. We then generated simulated behaviour using the empirical sequence of trials for that subject, preserving the goal sequence from that subjects’ data. Simulated trajectories were summarised with the same excess-steps calculation as the empirical data, with group mean across draw sets, and shaded regions showing the central 95% interval across replicated group means.

## Supplementary Figures

**Figure S1:**
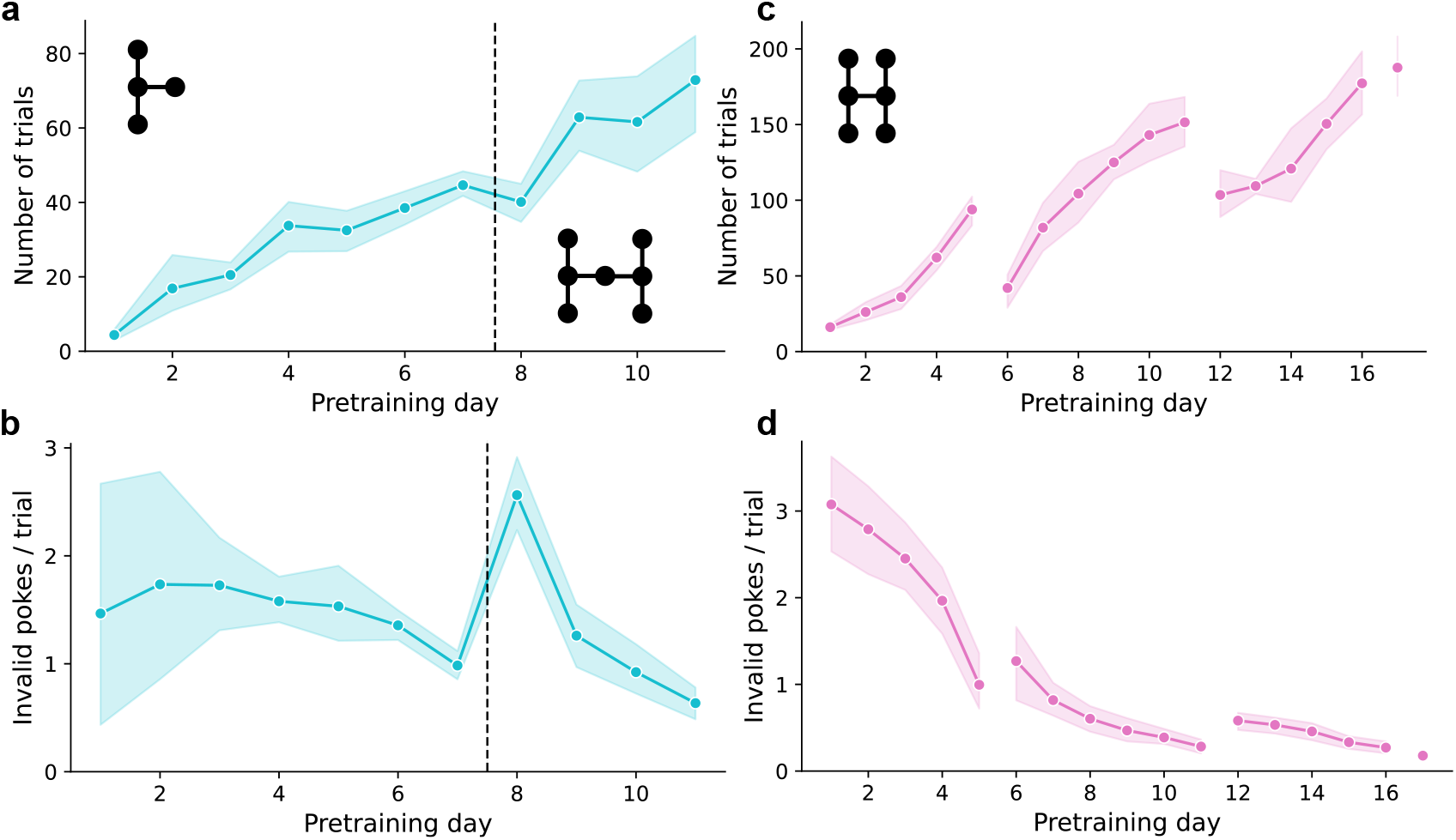
Pretraining behaviour. **a)** Number of trials per pretraining day in the pilot experiment. For the first 7 days of pretraining the maze configuration was a 4 node “T” shape, for the last 4 days of pretraining the maze configuration was a 7 node “H” (black inserts). **b)** Invalid pokes per trial - i.e. pokes in non-goal locations, across pretraining days in pilot experiment. **c)** Number of trials per pretraining day in the main experiment. For all days the maze configuration was a 6 node “H”. **d)** Pretraining invalid pokes per trial in main experiment. Lines show the mean across subjects, shaded regions show the 95% confidence interval. Lines are disconnected across non-consecutive calendar days.

**Figure S2:**
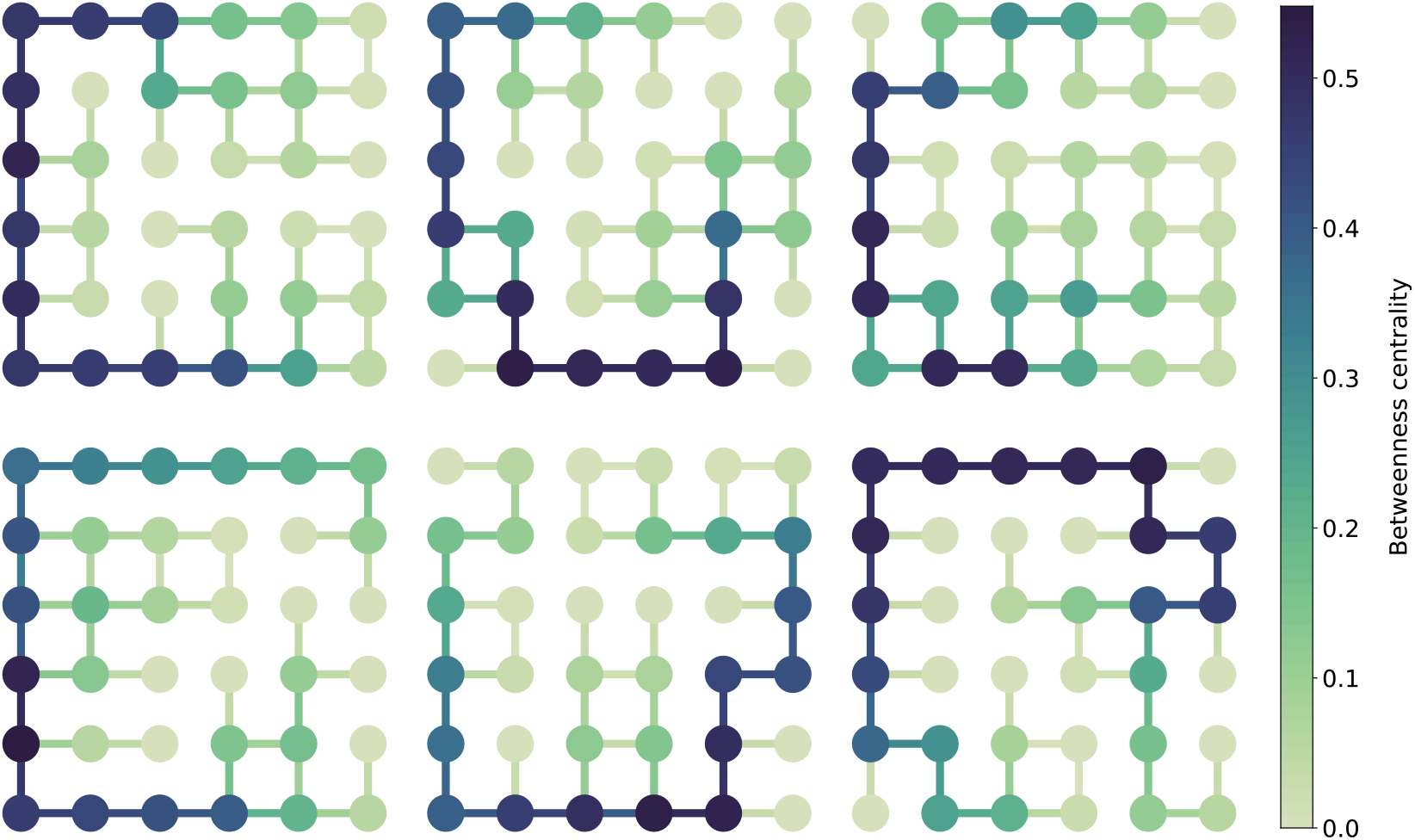
Optimised mazes: Top 6 candidate maze layouts generated by our maze search procedure when ranked by fraction of informative states in descending order. Note the prevalence of a prominent linear structure folded into the 6×6 grid.

**Figure S3:**
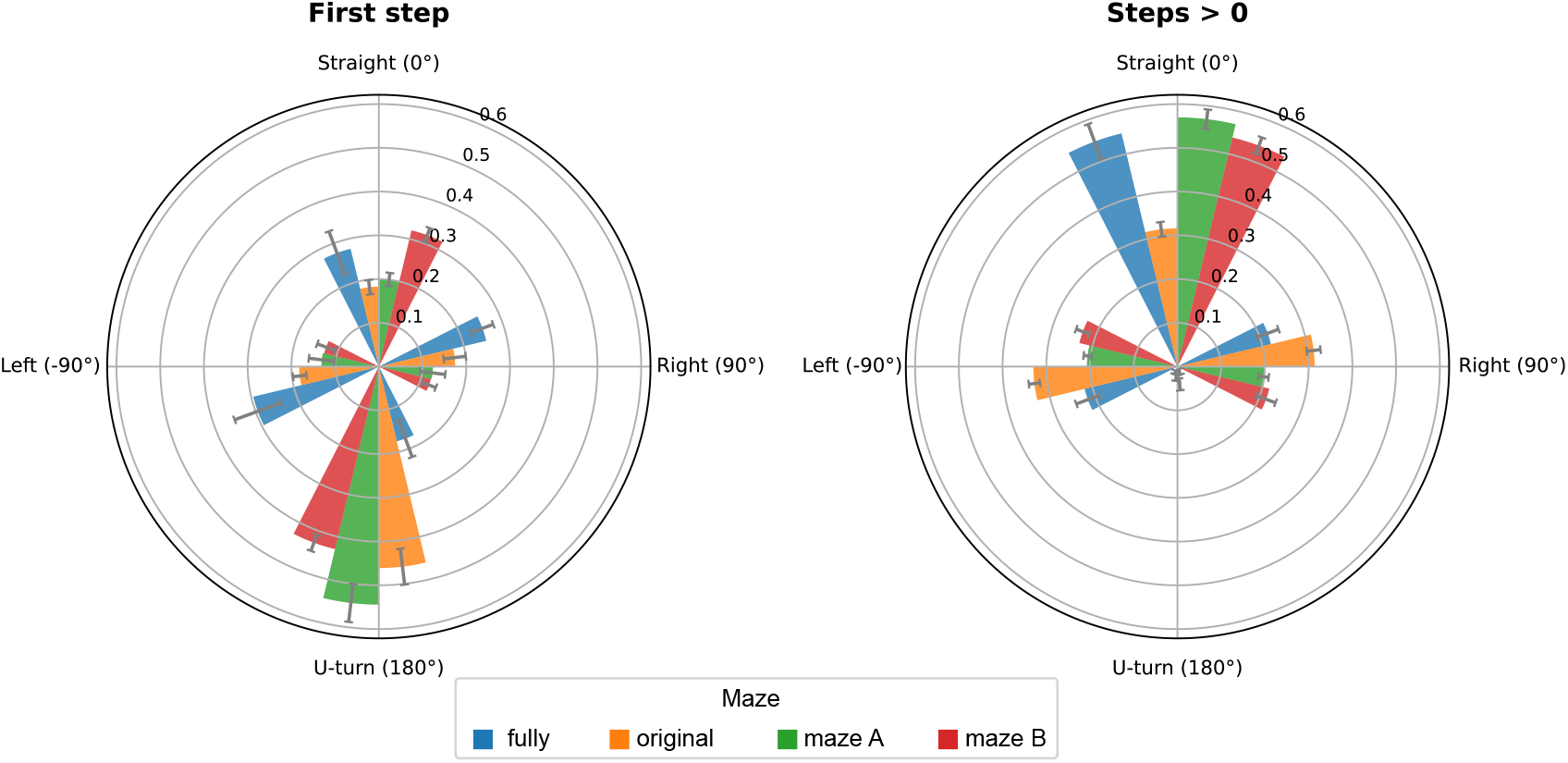
Egocentric action bias: Rate of taking the 4 possible egocentric actions; straight-on, left-turn, right-turn, u-turn, across the 4 mazes of experiment 2 (indicated by colour). Shown separately for the first step following goal cue onset (left panel) and for all subsequent steps during navigation to goal (right panel). The analysis was restricted to locations with more than one available action, excluding dead ends where the only possible movement is back along the previous path.

**Figure S4:**
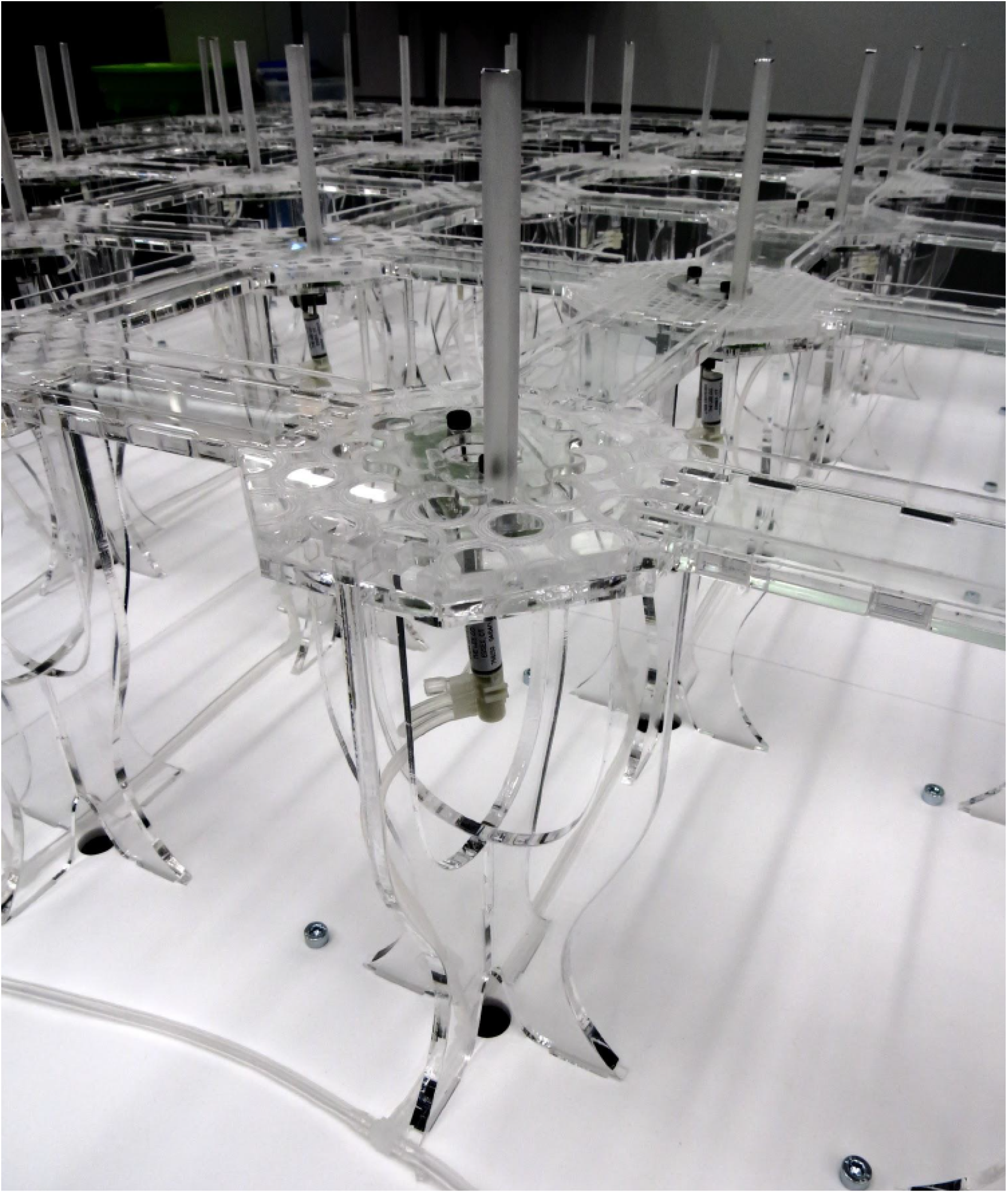
Tower and walkway construction. Photo showing close-up of a tower and connected walkways. The solenoid used to control reward delivery can be seen under the tower. An acrylic rod illuminated by the goal-cue LED sticks up vertically from the tower top.

## Supplementary Tables

**Table S1:**
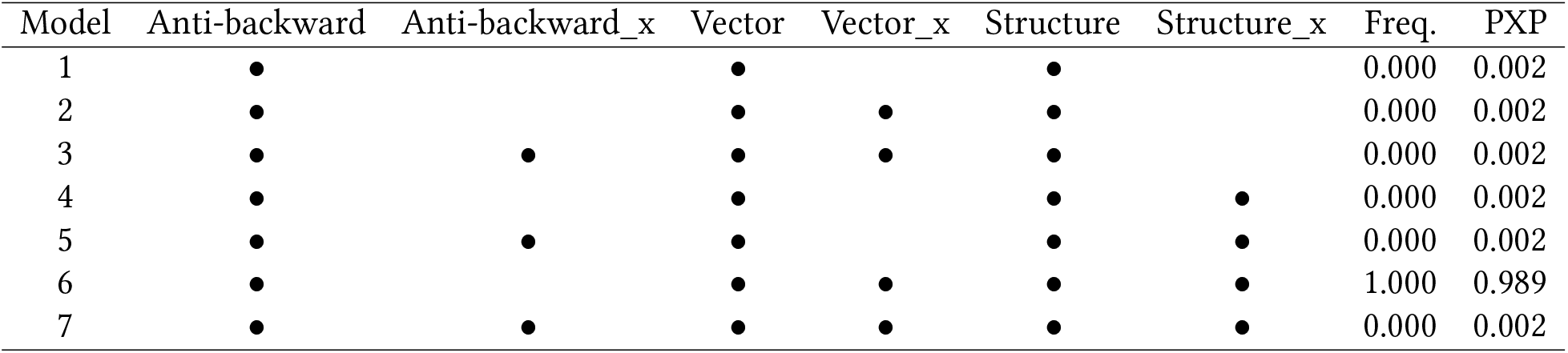
Model comparison summary. Bullets mark predictors included in each candidate model. ‘_x’ suffix indicates interaction with experience on maze, Freq. = estimated population model frequency, PXP = protected exceedance probability.

